# Feel the way with a vibrotactile compass: Does a navigational aid aid navigation?

**DOI:** 10.1101/122994

**Authors:** Steven M. Weisberg, Daniel Badgio, Anjan Chatterjee

## Abstract

Knowing where north is provides a navigator with invaluable information for learning and recalling a space, particularly in places with limited navigational cues, like complex indoor environments. Although north is effectively used by orienteers, pilots, and military personnel, very little is known about whether non-expert populations can or will use north to create an accurate representation of an indoor space. In the current study, we taught people two non-overlapping routes through a complex indoor environment, with which they were not familiar – a university hospital with few windows and several turns. Along one route, they wore a vibrotactile compass on their arm, which vibrated continuously indicating the direction of north. Along the other route, they were only told where north was at the start of the route. At the beginning, the end, and back at the beginning of each route, participants pointed to well-known landmarks in the surrounding city and campus (external landmarks), and newly-learned landmarks in the hospital (internal landmarks). We found improved performance with the compass only for external landmarks, driven by people’s use of the availability of north to orient these judgments. No such improved orientation occurred for the internal landmarks. These findings reveal the utility of vibrotactile compasses for learning new indoor spaces. We speculate that such cues help users map new spaces onto familiar spaces or to familiar reference frames.

---

Getting lost in complex indoor spaces, like hospitals, airports, or subways, can be dangerous, distressing, and costly. Unlike outdoor spaces, which typically offer long sightlines and distal landmarks, indoor spaces are often confined, undifferentiated, and labyrinthine. Individual building layouts can be hard to learn if knowledge about direction with respect to the larger world is not easily attained. Confusing layouts, signs, and an inability to see outdoor landmarks compound these problems. Even in places with adequate signs or sight of outdoor landmarks, successful navigation by blind people and people with visual impairments is a substantial problem (Schinazi, 2008; Schinazi, Thrash, & Chebat, 2016). In the current study, we investigate whether providing non-visual directional information aids sighted people in learning an unfamiliar, complex indoor space. Specifically, we tested people using a commercially-available vibrotactile compass, which vibrates in the direction of north. Unlike visual compasses, people with visual impairments can use vibrotactile compasses. A non-visual aid might also limit the effects of divided attention, which complicates the use of visual aids (Gardony, Brunyé, & Taylor, 2015).

We tested two competing hypotheses in the current study. These hypotheses stem from various theoretical frameworks of navigational ability (Wolbers & Hegarty, 2010).

1. General: People would have access to more spatial information, and thus improve generally. In this case, providing a specific spatial cue decreases the error in the sensory signals, increasing the fidelity of the resulting spatial representations. We would expect improved performance across all aspects of navigation behavior.
2. Specific: People would have access to a specific spatial cue, and thus improve on tasks for which that cue is immediately helpful. In this case, providing a specific spatial cue only improves access to specific information, but it does not create a more accurate general spatial representation.

Addressing these questions offers insights into how spatial information is acquired in general, and provides insight into how vibrotactile compasses might aid blind or visually impaired navigators.

Why might people get lost in complex indoor spaces? One reason is that place-based strategies used by humans to navigate are thwarted by indoor spaces. Research on humans and rats reveals two navigational strategies (Hartley, Maguire, Spiers, & Burgess, 2003; Marchette, Bakker, & Shelton, 2011; McDonald & White, 1994; Morris, Garrud, Rawlins, & O’Keefe, 1982; Munn, 1950; Packard & McGaugh, 1996; Restle, 1957; Tolman, Ritchie, & Kalish, 1946). First, a place-based strategy represents space as a cognitive map. A cognitive map is a flexible spatial representation, from which one can infer novel shortcuts. Second, a response strategy applies a stimulus-response approach to navigation. Choice points are identified and responses are recalled (e.g., turn left at the bank, then right at the big tree). The response strategy is relatively inflexible, and relies on associations between scenes and actions. In indoor environments, cues that support a place strategy, like distal landmarks and visual discriminability (Restle, 1957) are more likely to be absent. Instead, place strategies rely on path integration – tracking one's movement away from a starting location by attending to translations and rotations. Unfortunately, unlike rodents, people are poor path integrators (Foo, Duchon, Warren, & Tarr, 2007; Loomis et al., 1993). Thus, in indoor spaces, if people adhere to response-based strategies, they are less likely to gain spatial knowledge about the overall environment.

A second reason people easily get lost in complex indoor spaces could arise because of their use of preferred reference frames (Diwadkar & McNamara, 1997; McNamara, Rump, & Werner, 2003; Meilinger, Riecke, & Bülthoff, 2014; Meilinger, Frankenstein, Watanabe, Bülthoff, & Hölscher, 2015; Mou & McNamara, 2002; Mou, McNamara, & Zhang, 2013; Mou & Wang, 2015; Roskos-Ewoldsen, McNamara, Shelton, & Carr, 1998; Shelton & McNamara, 2004). A reference frame is a spatial representation in which objects or other representations of space are contained, or with respect to which they are ordered, oriented, located, or thought to move; a preferred reference frame refers to an individual's privileged reference frame, according to which a spatial layout is most easily recalled. In a global reference frame, properties do not change across different areas (e.g., global north). In familiar, large-scale (i.e., city-sized) spaces, reference frames can be aligned with north (Frankenstein, Mohler, Bülthoff, & Meilinger, 2011), or a salient organizing feature (like a street through a campus; Marchette, Yerramsetti, Burns, & Shelton, 2011; Yerramsetti, Marchette, & Shelton, 2013). Indoor environments, however, provide less access to global reference frames. In the absence of global cues (like distal landmarks), people typically organize space into several local reference frames, instead of one global reference frame (Meilinger et al., 2014). In the extreme, using local reference frames means each segment of a route is disconnected from the last, and results in a 1-dimensional representation of space (i.e., an ordered sequence of places, with no 2-dimensional spatial relations specified; Ishikawa & Montello, 2006). Being able to integrate across areas of a learned route to take novel shortcuts, for example, requires spatial knowledge about how to join local reference frames to each other to form a global reference frame – a difficult and demanding cognitive process (Weisberg & Newcombe, 2016; Weisberg, Schinazi, Newcombe, Shipley, & Epstein, 2014).

If people directly sense a global spatial cue, like a cardinal direction, they might map an unfamiliar indoor space onto the larger external environment, and construct a more accurate spatial representation of this indoor space. In this way, place-based strategies and global reference frames become more useful in an unfamiliar indoor space. Vibrotactile compasses in particular can convey global north, with some success. König and colleagues (Kärcher, Fenzlaff, Hartmann, Nagel, & König, 2012; Kaspar, König, Schwandt, & König, 2014; König et al., 2016) developed a feelSpace belt, which provides tactile information about true north. Their work illustrates that, with extensive training (seven weeks), a vibrotactile compass improves basic homing tasks in sighted individuals (König et al., 2016). The focus of the current study is on how such information improves (or does not improve) specific and general aspects of navigation. That is, we do not know if such directional information is incorporated into different spatial representations, particularly in complex indoor spaces. Moreover, we wished to learn if this directional information could be used to learn a new environment with only rudimentary use of the device.

## Method

### Participants

We recruited 52 participants from the University of Pennsylvania using an online recruitment tool specific to that university. We excluded data from four participants because they were highly familiar with the hospital (having spent over 30 hours in the environment within the last year). Of the remaining subjects, 18 had been in the hospital, but the maximum amount of time spent by any one subject was 4 hours, with an average of 1.97 hours. The resulting sample consisted of 48 participants (27 identifying as female). Sixteen participants self-reported as Asian, eight as African-American or Black, five as Hispanic or Latino, two as Other, and 17 as Caucasian or White. Participants' average age was 21.73 years (SD = 3.43). This research was approved by the institutional review board of the University of Pennsylvania.

### Experimental settings and materials

#### Testing rooms

We used a small testing room to greet the participant, obtain informed consent, and have the participant fill out the initial questionnaires. The larger testing room was either the first or third author's office, and provided enough space for the participant to be blindfolded and disoriented without bumping into furniture or the walls.

#### Hospital

We chose two routes through the main corridors of a large, complex university hospital (Fig. 1 and Fig. 2A). In general, this part of the hospital has non-descript hallways and lacks windows or views of the outside. Occasional signs mention a street exit (e.g., to Spruce Street), but the majority of the signs consists of hospital building names and colors. The routes were on different floors so that they did not overlap. Each route took 3-4 minutes to walk, and contained three internal landmarks, or objects, about which the participant was instructed to learn certain details (see Measures and Procedure, below).

**Figure 1.**
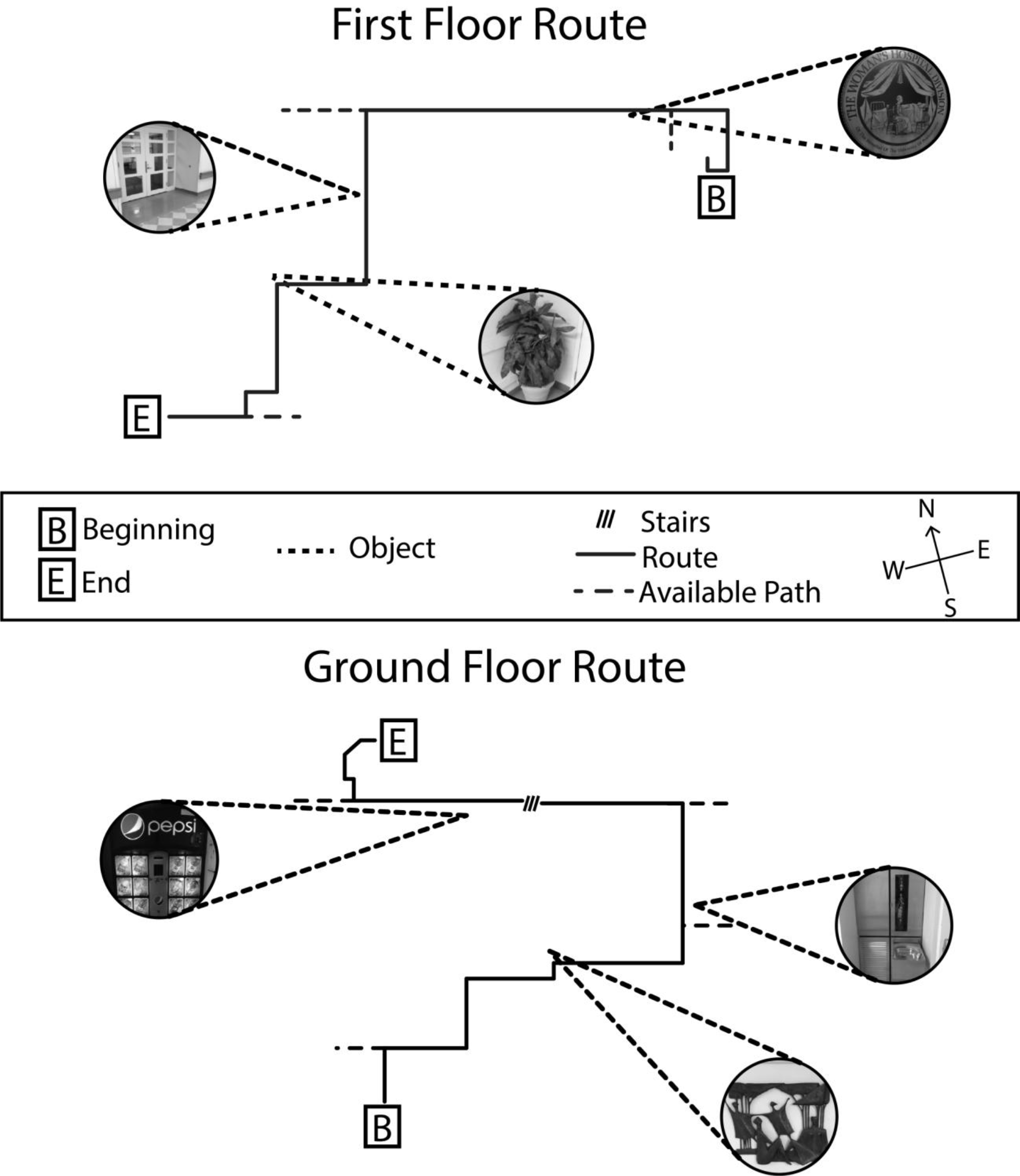
Schematic maps of the first floor and ground floor routes. Images depict internal landmarks and colored lines indicate their positions along the route. Dotted lines indicate available alternative pathways at which participants could have made wrong turns. Other turns along the route were not decision points. As shown, the routes are aligned with north, but are not vertically aligned with each other (i.e., the first floor route was actually mostly northeast of the ground floor route). (For a color version of this figure, see the online version of the article.)

**Figure 2.**
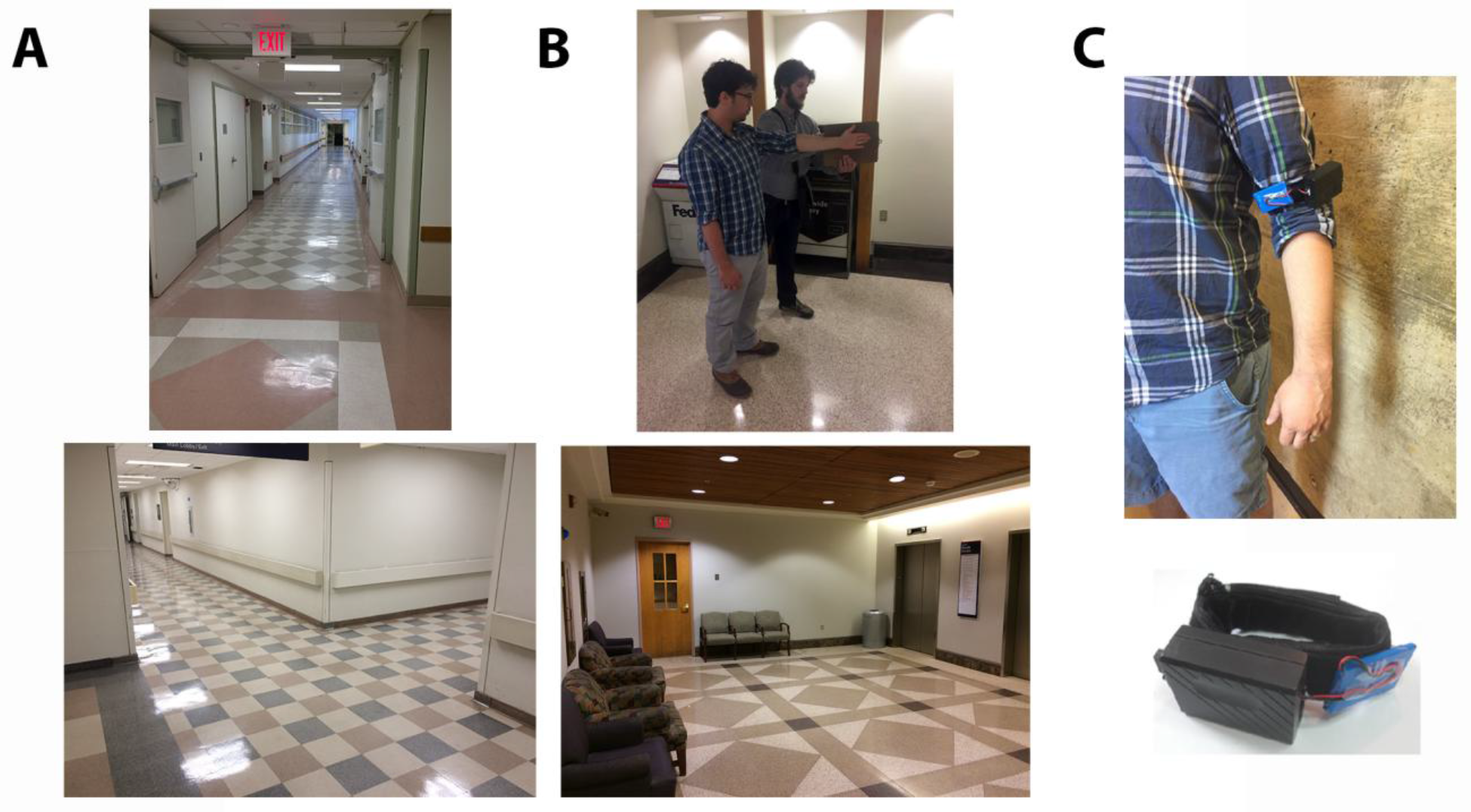
Photographs of typical hallways (A), and endpoints where the pointing test, sketch map task, and post-test questionnaire were administered (B) from along the first floor route (top, A and B) and ground floor route (bottom, A and B). On the bottom right, the pointing task and measurement is being demonstrated by two researchers. The participant (foreground) points using his arm, while the experiment lines a clipboard up with the direction and measures it with the digital compass (not shown). The digital compass was not visible to the participant. Photographs of the vibrotactile compass (C) show how it was worn on the arm (top) and the housing (black box), battery (blue pad) and motor enclosure (black velvet). (For a color version of this figure, see the online version of the article.)

The starting and ending locations were in areas removed from heavy foot traffic (e.g., a small shipping alcove connected to a lesser-used section of hallway). These locations were also the points at which the participant would complete the pointing task (see Figure 1). The ground floor route consisted of ten turns (three choice points). The first floor route consisted of nine turns (four choice points).

#### Vibrotactile compass

We used a vibrotactile compass (Fig. 2B; NorthPaw 2.0; hereafter referred to as the compass), which was purchased pre-assembled from an online vendor (www.sensebridge.net). The compass consists of eight oscillating motors arranged approximately one inch apart around a cloth strap, and an Arduino circuit board and battery assembly. The cloth strap was wrapped around the bicep of the participant’s non-dominant arm. The eight motors were programmed to vibrate based on whichever one was facing closest to true north, as measured by a compass sensor wired into the Arduino circuit board. Thus, as the participant moved and rotated around, the motor that was closest to north would vibrate. Because one motor is always the closest to north, exactly one motor vibrated the entire time the compass remained on. The experimenter explained this to the participant, and demonstrated the compass' functionality.

#### Pointing measurement compass

For the majority of participants^1^, to measure the participant's pointing judgments, we used a digital compass (purchased from Amazon.com https://www.amazon.com/Pyle-Sports-PSHTM24-Handheld-Chronograph/dp/B005G2SH8Y). We measured pointing direction by placing a clipboard along the direction of the participant's arm as they pointed, then aligning the compass with the clipboard, and recording the direction to the nearest degree. This procedure was used so that the participant could not view the direction they were pointing.

### Measures

#### Paper and pencil measures

##### Demographics

Demographics and self-report data were collected on age, ethnicity, sex, gender, education (in years), sexual orientation, English as a first language, and whether the subject was right-handed.

##### Familiarity with hospital questionnaire

This questionnaire consisted of three questions: 1) Have you been inside the [hospital] before for any reason? 2) If yes, how long ago? 3) Approximately how many hours have you spent inside the hospital?

##### Santa Barbara Sense of Direction Scale

(SBSOD; Hegarty, Richardson, Montello, Lovelace, & Subbiah, 2002)). This self-report measure of navigation ability consists of fifteen 7-point Likert-scale items such as "I am very good at giving directions," and "I very easily get lost in a new city." The average score for each participant has been shown to correlate highly with performance on behavioral navigation tasks in real and virtual environments (Hegarty et al., 2002; Weisberg et al., 2014), and with individual differences in neural structure and function (e.g., Epstein, Higgins, & Thompson-Schill, 2005; Schinazi, Nardi, Newcombe, Shipley, & Epstein, 2013).

##### Spatial Cognition, Navigation, and Experience Questionnaire

(SCNEQ). We developed this 7-question self-report survey to measure the extent to which each participant was accustomed to navigating similar urban environments. For example, two items were “I lived much of my life in a rural setting” and “I am used to the layout of a city whose roads are arranged in a grid.” See Supplementary Methods for the full versions of all questionnaires.

#### Navigation performance measures

##### Compass pre-test

The experimenter introduced the participant to the compass, explained and demonstrated how the mechanism worked, and described how it could be used to help maintain an orientation. The participant then completed a pre-test pointing task. The purpose of this pre-testing task was to familiarize them with the procedures required during the main experiment. Across participants, we counter-balanced the order of the pre-test. Half of the participants first completed the pointing without the compass, and the other half first completed the pointing with the compass. The procedure was the same for both. The experimenter pointed in a pre-determined random direction (between 0-359, with 0 as true north), placed a blindfold on the participant, slowly spun them around clockwise and counterclockwise for 20-30 seconds, then asked the participant to point in what they thought was their original facing direction and recorded the judgment. This process was repeated twice, disorienting in the same way to record a second and third pointing judgment for the same direction. The participant remained blindfolded continuously throughout the duration of all three trials. The entire procedure was then executed with a new direction and with the compass turned on (or off, depending on what the participant had already completed).

##### Pointing judgments

Along each hospital route, participants completed a pointing task three times on each route: at the beginning before route learning, at the end of the route learning, and again at the beginning after route navigating. All three pointing sessions included the four cardinal directions (North, East, South, and West) and eight familiar external landmarks. Four of these landmarks were located on the university campus (External-Campus), and four were located in Philadelphia (External-City). The second and third pointing sessions included pointing judgments for three internal landmarks, which were located along that route. Participants pointed directly at each internal landmark (Internal-Location), and pointed in the direction they were facing when they viewed that landmark (Internal-Facing). The third pointing session of the participant’s second route also included judgments of the internal landmarks of the first route.

##### Route reversal

After learning the route in one direction, the participant navigated back to the beginning. If the participant made a navigational mistake (e.g., by making a wrong turn), the experimenter corrected it immediately. Along the way, they indicated and named the internal landmarks, and any apparent potential short cuts back to the start (e.g., doorway, hallway), without actually taking them. Data were collected by the experimenter on numbers of navigational errors made, numbers of correctly indicated short cuts, and numbers of missed internal landmarks.

##### Sketch map

Following the third pointing task on each route, the participant sketched a map of the route they took. The maps were required to include the path of the route itself, the internal landmarks, and an arrow indicating north.

##### Post-route questionnaire

After the sketch map, participants answered four de-briefing questions about each route. 1) Explain how you tried to learn the route around the hospital and the locations of the objects. 2) Describe the route you took (in words). 3) What strategy did you use to perform the pointing task? 4) (Compass route only) How did the device help (or hurt) you in learning the space and re-walking the route?

### Procedure

Participants met the experimenter in a small testing room to provide informed consent and fill out four questionnaires (demographics, familiarity with the hospital, SBSOD, SCNEQ). The experimenter led the participant to a larger room for the compass familiarization and pre-test. The experimenter then took the participant from the meeting place to the hospital. The experimenter took the same path every time to enter the hospital.

At the hospital, the route order and compass-use order were counter-balanced between participants such that all four possibilities were equally represented in the sample. The experimenter led the participant to the beginning of her first route, and explained the general nature of the experiment (see supplemental methods for all experimenter scripts).

Each route consisted of the same tasks in the same order. First, the experimenter pointed in the direction of north, and told the participant that she could use that information however she saw fit. The participant then pointed to the cardinal directions and all external landmarks.

The experimenter then outlined the participant's tasks along the route – to learn the order, locations, and facing orientations of three objects, and to learn the layout of the route with respect to the hospital and the broader external environment. For the Compass route, the experimenter strapped the compass on, and instructed the participant to use it as a helpful tool in maintaining her orientation throughout the environment. The experimenter then led the participant along the route, stopping at each object to name it. The experimenter also positioned the participant in front of each internal landmark in a consistent location, indicating the direction the participant was facing as the facing orientation to that internal landmark. At the end of the route, the participant performed the pointing task again, beginning with the cardinal directions and external landmarks, and including the internal landmark locations and facing orientations. The participant also pointed back to the start of the route.

The participant then led the way back from the end of the route to the beginning (the Route Reversal task), indicating each internal landmark along the way. Upon returning to the beginning, the participant performed all pointing judgments a third time, this time including pointing to the end of the route. If this was the second of the two routes, the participant then pointed to the internal landmark locations and facing orientations of the previous route. Finally, the participant drew a sketch map of the route, and answered the post-test questionnaire. If this was the first of the two routes, the experimenter would walk the participant over to the beginning of the second route and repeat the foregoing procedure.

## Results

Our primary interest was in differences when participants were wearing the compass and not. We also did not predict, and were not interested in interactions with route order or order of compass use. We thus collapsed data across subjects for route order and order of compass use. To calculate the error for each pointing judgment we subtracted the angular distance between the correct answer and the participant's response, then corrected the difference to be less than 180°. Chance performance would average 90° error. For repeated-measures *t*-tests, we used the correction to Cohen's *d* suggested by Morris & DeShon (2002), which accounts for correlations between factors of the dependent variable. For repeated-measures effect sizes, we use generalized *eta squared* (Bakeman, 2005) provided by the ezAnova package in R, 3.0. For the sketch maps, we scanned and uploaded the hand-drawn maps, then calculated coordinates for the three landmarks, the starting point, and the finishing point, using the Gardony Map Drawing Analyzer (GMDA; Gardony, Taylor, & Brunyé, 2016).

### Compass Pre-test

Participants pointed to the indicated direction better than chance both with the compass, *t*(47) = 6.14, *p* < .0000001, *d* = 0.89, and without the compass, *t*(47) = 2.89, *p* < .006, *d* = 0.42. We compared performances using a within-subjects t-test on the pre-test and found that participants were significantly more accurate with the compass (*M* = 58.78, *SD* = 35.24) than without the compass (*M* = 76.74, *SD* = 31.81), *t*(47) = 2.81, *p* = .007, *d* = 0.41.

### Route-reversal

Overall, participants made fewer navigation errors when recreating the route with the compass (total errors = 27) than without the compass (total errors = 35). These did not differ significantly: *t*(47) = 1.31, *p* =.20, *d* = 0.19. Participants also did not make more mistakes naming the objects along the route, were not more likely to indicate the wrong direction for the stairs, and did not indicate more shortcuts, all *p*'s > .43. The additional input from the compass did not improve the participants' ability to effectively re-create the route, although we note that most participants were at a ceiling performance for this aspect of navigation.

### Pointing Test Error

We assessed differences in pointing separately for each judgment type (cardinal directions, external landmarks, internal locations, and internal orientations) using a 2-factor (Compass x Route position) within-subjects ANOVA. Compass had two levels – with or without the compass. Route position also had two levels – End, and Beginning 2. We excluded judgments from the Beginning because a) participants were shown where north was during that pointing session, and b) no judgments were made to Internal Landmarks. Furthermore, no differences were observed between the two Beginning sessions, and the pattern of results is similar for the Cardinal Directions and External Landmarks when the two Beginning sessions are averaged together as when only Beginning 2 was used. For simplicity and brevity, we thus exclude Beginning judgments.

#### Cardinal directions

We obtained a significant interaction between compass and route position, *F*(1,47) = 9.65, *p* = .003, 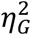 = 0.04. As predicted, follow-up contrasts revealed that pointing at the End was significantly better when participants wore the compass (*M* = 25.06, *SD* = 25.06) than when they did not (*M* = 52.33, *SD* = 54.22), *t*(47) = 3.46, *p* = .001, *d* = 0.55. Participants performed equally well at Beginning 2 with the compass (*M* = 14.75, *SD* = 10.32) as without the compass (*M* = 16.75, *SD* = 8.56) *t*(47) = 1.58, *p* = .12, *d* = 0.23. The interaction resulted in a significant main effect of wearing the compass, *F*(1,47) = 13.95, *p* < .001, 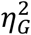 = 0.06, such that participants were more accurate when wearing the compass overall (*M* = 19.91, *SD* = 13.90) than without the compass (*M* = 34.54, *SD* = 27.50), and in a main effect of route position, *F*(1,47) = 23.97, *p* < .001, 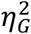 = 0.13, such that pointing to cardinal directions at Beginning 2 (*M* = 15.75, *SD* = 8.40) was significantly more accurate compared to the End (*M* = 38.70, *SD* = 32.21). Thus, participants became disoriented with respect to north after walking the route, and this disorientation was mitigated by use of the compass.

#### External Landmarks

We obtained a significant interaction between compass and route position for external landmark judgments, *F*(1,47) = 4.47, *p* = .04, 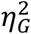 = 0.01. As predicted, follow-up contrasts revealed that pointing at the End was significantly better when participants wore the compass (*M* = 49.41, *SD* = 28.50) than when they did not (*M* = 61.58, *SD* = 39.69), *t*(47) = 2.34, *p* = .02, *d* = 0.35). There was no difference at Beginning 2 with the compass (*M* = 44.98, *SD* = 25.61) compared to without the compass (*M* = 44.89, *SD* = 27.87), *t*(47) = 0.04, *p* = .97, *d* = 0.005. This is not surprising because at the Beginning of both routes, participants were shown where north was, and could have used this information to reorient themselves. The interaction between compass and route position resulted in a significant main effect of wearing the compass, *F*(1,47) = 4.66, *p* = .036, 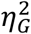 = 0.01, such that participants were more accurate with the compass (*M* = 47.19, *SD* = 24.34), than without the compass (*M* = 53.24, *SD* = 28.79). The interaction also resulted in a main effect of route position, *F*(1,47) = 9.31, *p* = .004, 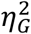 = 0.03, such that pointing to cardinal directions at the Beginning 2 (*M* = 44.94, *SD* = 25.51) was significantly more accurate compared to the End (*M* = 55.49, *SD* = 29.49). Thus, as with cardinal directions, participants lost their orientation with respect to the locations of external landmarks after walking the route, and the disorientation was mitigated by the use of the compass.

#### Internal Landmarks - Location

We did not obtain significant effects for the compass or route position, nor did we obtain an interaction. Participants were no better at pointing to the locations of the internal landmarks while wearing the compass at either the End (*M* = 49.59, *SD* = 33.95) or at Beginning 2 (*M* = 58.64, *SD* = 38.78) compared to when they were not wearing the compass at the End (*M* = 51.35, *SD* = 35.99) or at Beginning 2 (*M* = 55.72, *SD* = 39.19), all *p*'s > .12. Wearing the compass did not help pointing to the locations of internal landmarks.

#### Internal Landmarks - Orientation

We did not obtain significant effects for the compass or route position, nor did we obtain an interaction. Participants were no better at pointing to the orientations of the internal landmarks while wearing the Compass at either the End (*M* = 42.96, *SD* = 33.39) or at Beginning 2 (*M* = 47.92, *SD* = 33.95) compared to when they were not wearing the Compass at the End (*M* = 56.46, *SD* = 47.24) or at Beginning 2 (*M* = 46.94, *SD* = 44.28), all *p*'s > .13. Again, the compass did not help pointing to the orientations of internal landmarks.

### Pointing Reference Frames

By analyzing pointing errors, we found that participants were disoriented with respect to cardinal directions and external landmarks at the End of the route when they were not wearing the compass. From the pointing error analysis, we did not see improvement between compass and non-compass routes on the internal landmark orientation or location judgments. This result could be because participants were not able to use the compass to improve these judgments, despite attempting to align the representation of internal landmarks with north. Therefore, we also wanted to know whether the reference frame participants used for the internal landmarks mapped onto their reference frame for the external landmarks. If so, we would expect pointing judgments within a session to be internally consistent with each other. This would mean that each pointing judgment might have a large absolute error, but that overall, the pointing judgments might be internally consistent but with an orientation deviated from true north. We tested this idea by finding the Best Fit Angle for each set of pointing judgments (see top panel of Figure 3).

**Figure 3.**
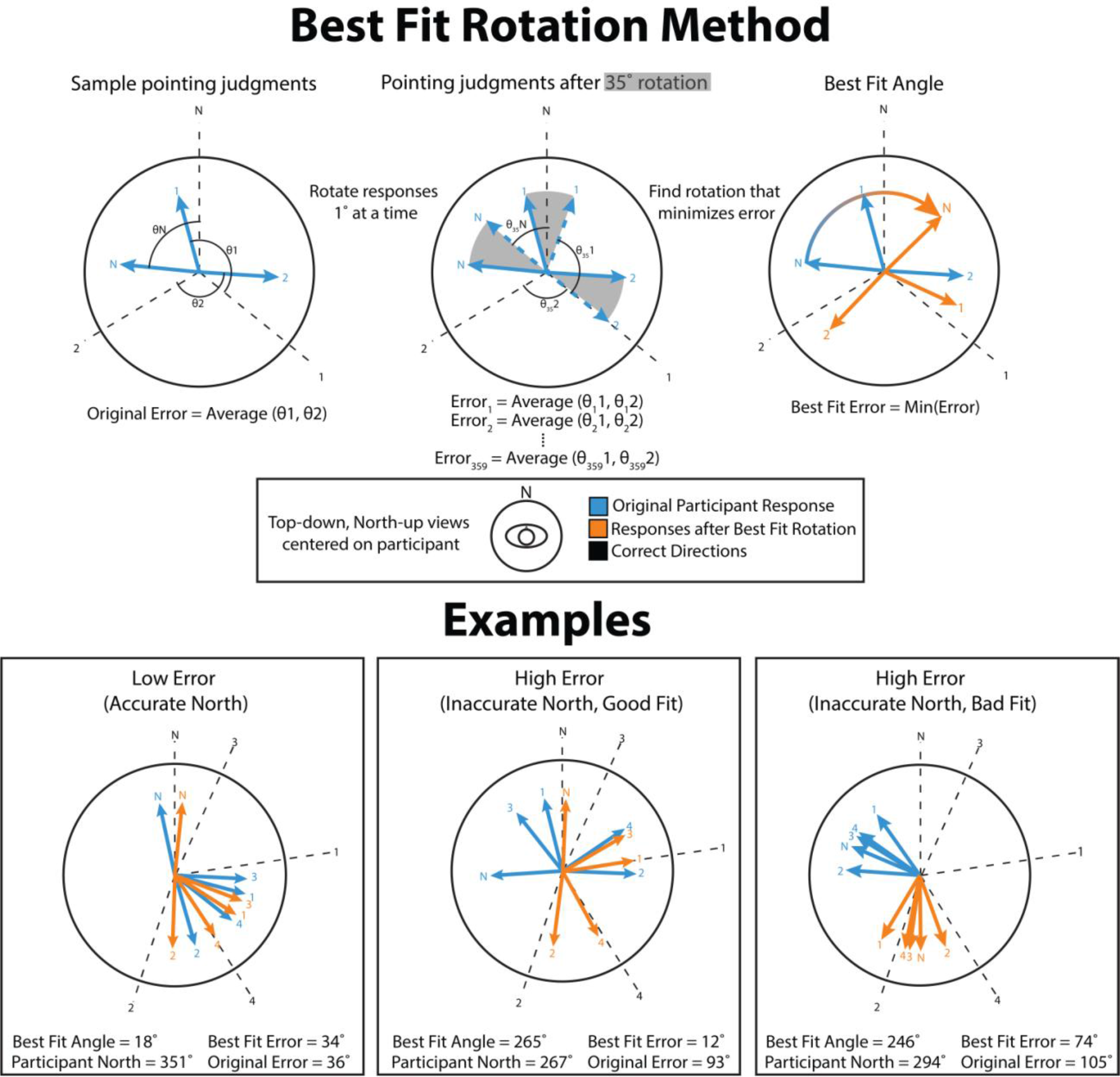
Illustration of the best fit rotation method (top) and examples of schematized participant data (bottom). The best fit rotation method seeks to find the optimal rotation of a set of pointing judgments to minimize their combined error. To do so, pointing judgments are collected (top left), and the angle between each judgment and the correct direction is measured (correcting for angles greater than 180°). Next, the set of pointing judgments is rotated, 1° at a time, and the error is recalculated (top center). This is done for all 360°. The angle for which the minimum error is recorded is the Best Fit Angle (top right). The examples (bottom) show the three categories of outcomes for a set of pointing judgments. If there is low error to begin with (bottom left), the best fit method will usually result in a small rotation, with little decrease in error. If there is high error, there are two possibilities. One possibility is that pointing judgments are internally consistent, but not aligned with the actual answers (bottom center). In this case, the best fit rotation will likely yield an answer that is close to where the participant thought north was, and will substantially decrease the error. The second possibility is that pointing judgments are internally inconsistent (bottom right). In this case, the best fit rotation will likely yield an answer that is not close to where the participant thought north was, and will not substantially reduce the error. If participants are disoriented without the compass, we should see many inaccurate north, Good Fit solutions. With the compass, we should see a greater number of low error overall solutions.

This approach allows us to infer the orientation of a set of pointing judgments. Note that we could use the direction that the participant pointed to when asked where north was. But this approach would not allow us to find an optimal solution for internal landmarks, which could be aligned with a local reference frame, or otherwise misaligned with where the participant thought north was, but still internally consistent.

The lower middle panel of Figure 3 illustrates an example wherein a participant believes that north is actually west, and was mapping other judgments according to that misaligned orientation. Each pointing judgment is thus off by approximately 90°. When the pointing judgments are rotated to minimize the error, true north lines up with Participant-north, suggesting that the participant's pointing judgments were consistent with where he thought north was. Applying the Best Fit Method allows us to see whether participants use self-consistent reference frames for external and internal landmarks.

To find the Best Fit Angle, we rotated each participant's pointing judgments one degree at a time and calculated the absolute error. For each set of pointing judgments, we recorded the angle at which the minimum absolute error was achieved (henceforth the Best Fit Angle). We did this separately for the external landmarks and internal landmarks to obtain separate Best Fit Angles for each set of judgments. Because participants were told where north was at the Beginning (and likely recalled this information at Beginning 2), we only used judgments from the End. All circular statistics were calculated using the CircStat package in Matlab (Berens, 2009).

Error should be reduced overall for both routes, because finding the Best Fit Angle also minimizes error from noise. However, if participants pointed to landmarks based on a consistent frame, we would expect more error reduction for pointing without the compass compared to with the compass route. Consider the three examples in the bottom panel of Figure 3. In the example on the left, the participant knows where true north is, and has relatively accurate judgments for the landmarks. Rotating by the Best Fit Angle reduces the error by a very small amount. In the example in the middle, the participant does not know where true north is. He also does not know where the landmarks are with respect to north or with respect to each other. Thus, the Best Fit Angle does not line up with Participant-north. In this case, Best Fit Angle and Participant-north would be uncorrelated. Finally, in the example on the right, the participant thought north was actually west, and his landmark pointing judgments aligned with this "west-is-north" reference frame. In this case, the participant's Best Fit Angle is close to 90° (where 0° is north), which is also the angular difference between west and north. We expected this pattern to be common for the external landmarks when participants were not wearing the compass, because participants might become disoriented with respect to north but use their own sense of north to guide pointing to the external landmarks. We were most interested in the internal landmarks tasks. If participants used the compass to place and orient internal landmarks in terms of north, we expected the Best Fit Angle to substantially reduce the error more when participants did not wear the compass, and for the Best Fit Angle to correlate with Participant-north.

#### External Landmarks

For the external landmarks, one-sample *t*-tests revealed that rotating the pointing judgments, as expected, improved accuracy with the compass, *t*(47) = 4.24, *p* < .001, *d* = 1.24, and without the compass, *t*(47) = 4.95, *p* < .001, *d* = 1.44 (See Figure 4). However, the improvement without the compass was greater (*Mean Error Reduction* = 26.08, *SD* = 36.52) than with the compass (*Mean Error Reduction* = 13.88, *SD* = 22.68), *t*(47) = 2.36, *p* = .02, *d* = 0.36. The resulting Best Fit Error was the same with the compass, (*M* = 35.51, *SD* = 15.37) and without the compass (*M* = 35.51, *SD* = 17.73), *t*(47) = 0.001, *p* = .99, *d* < 0.01, confirming that despite being disoriented to true north participants remained consistent in their pointing based on an internal reference frame.

**Figure 4.**
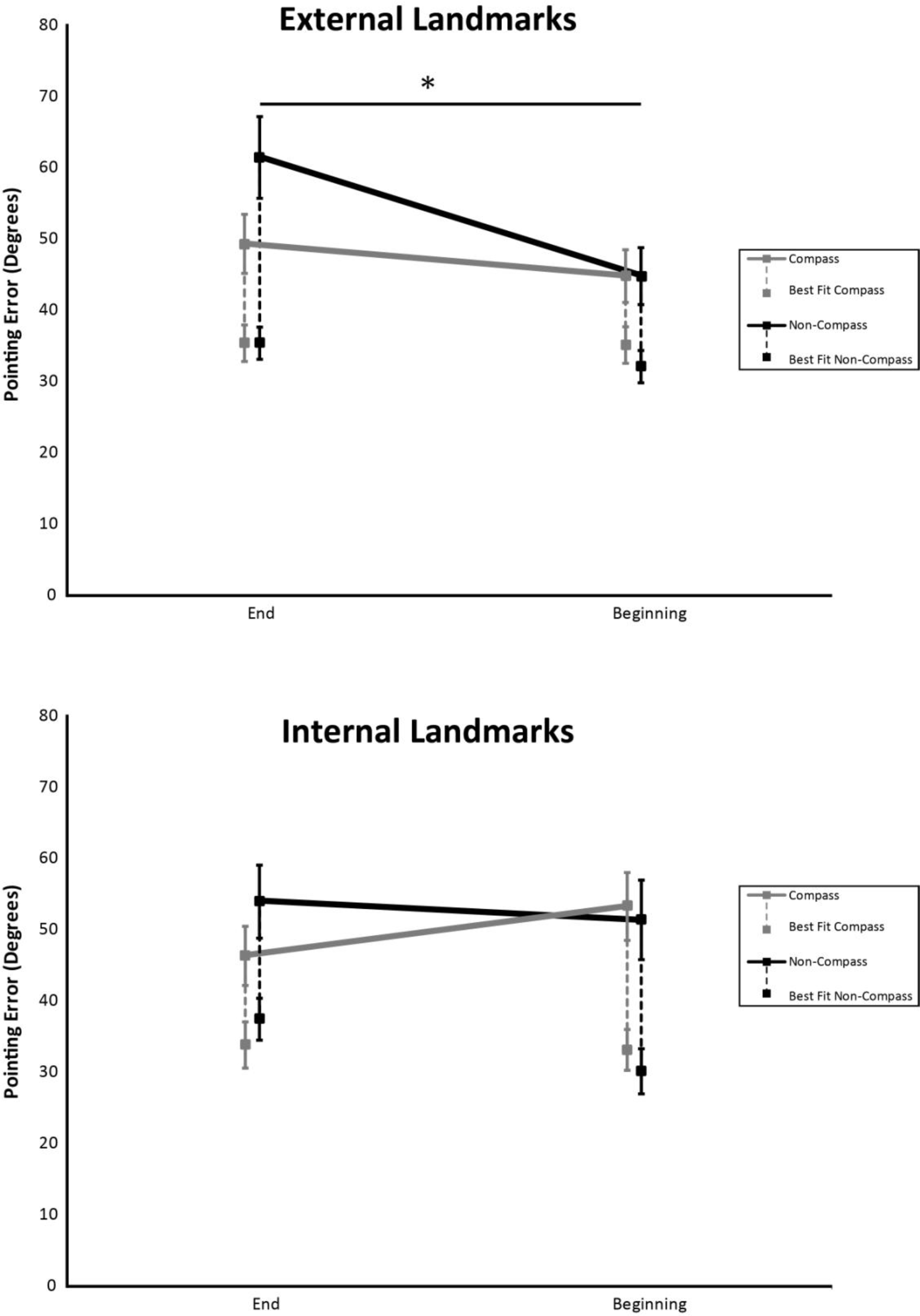
The pointing error for External Landmarks (above) and Internal Landmarks (below; Locations and Orientations). Unadjusted errors are higher, and connected by solid lines. Best fit errors are connected below with dotted lines. The asterisk highlights the significant interaction between Compass and Non-Compass pointing error between the End (where participants were not explicitly told where north was) and back at the Beginning (where they had previously been told where north was). Adjusting pointing error with the Best Fit method (Figure 2) eliminates this interaction. Such a pattern is not evident for the Internal Landmarks, where errors are relatively high at both pointing locations and with or without the compass.

To determine whether the external landmark Best Fit Angle correlated with the direction which the participant judged as north, we obtained the circular correlation between the Best Fit Angle with Participant-north. The correlation was significant without the compass, *r*(48) = .61, *p* < .001, and marginally significant with the compass, *r*(48) = .33, *p* = .066. This suggests that the Best Fit Angle for the external landmarks brought participants' pointing judgments into alignment with where they thought north was.

#### Internal Landmarks

For the internal landmarks, we combined location and orientation judgments.^2^ One-sample *t*-tests revealed that rotating the pointing judgments resulted in improved accuracy with the compass, *t*(47) = 5.69, *p* < .001, *d* = 1.66, and without the compass, *t*(47) = 6.83, *p* < .001, *d* = 1.99 (See Figure 4). Unlike for the external landmarks, the non-Compass route (*Mean Error Reduction* = 16.50, *SD* = 25.22) was not significantly different from the Compass route (*Mean Error Reduction* = 12.50, *SD* = 15.22), *t*(47) = 0.35, *p* = .94, *d* = 0.14. The resulting Best-fit error was the same for the Compass route, (*M* = 33.78, *SD* = 22.12) and the non-Compass route (*M* = 37.42, *SD* = 20.67), *t*(47) = 1.06, *p* = .29, *d* = 0.15. This analysis confirms our basic finding that the compass did not help people in pointing accurately to the location of internal landmarks.

Again, we wondered whether Best-Fit Angle correlated with where the participant thought north was. Significant correlations would be evidence that participants are using participant north to guide their estimation of the internal landmark locations and orientations. Using circular correlations, the Best Fit Angle did not correlate with Participant-north without the compass, *r*(48) = .04, *p* = .77, or with the compass, *r*(48) = -.12, *p* = .41.

### Sketch Maps

We discarded the data of two participants who did not provide all the requested data for one of their maps, resulting in N=46. We wanted to know whether knowledge of the configuration of the internal landmarks in their sketches, including the starting point and ending point, improved when participants wore the compass. To measure configuration, we calculated a bidimensional regression for each map. Bidimensional regression (Tobler, 1994) corrects for deviations in scale, translation, and rotation, then provides a measure of similarity (here we use *r*^2^, which can be interpreted as the proportion of variance explained) between the actual configuration of points, and a reconstruction of those points. The configuration of internal landmarks was not significantly different with the compass (*M* = .73, *SD* = .25) than without the compass (*M* = .69, *SD* = .26), *t*(45) = 0.89, *p* = .38, *d* = 0.14. Data from the sketch maps corroborated the finding from the Internal Landmarks Location pointing judgments: the compass did not improve participants' ability to represent the locations of these landmarks.

### Verbal Descriptions

In the verbal descriptions participants provided for each route, we coded for the amount of relative (right, left, ahead) and absolute (north, east, south, or west) spatial language used. The vibrotactile compass did not increase use of absolute spatial language in describing the routes. Participants used a similar number of absolute spatial terms in the Compass (*M* = 1.75, SD = 3.31) and Non-Compass (*M* = 1.98, *SD* = 3.34) conditions, *t*(48) = 0.73, *p* = .47, *d* = 0.11. Participants also used a similar number of relative spatial terms in the Compass (*M* = 7.06, *SD* = 5.54) and Non-Compass (*M* = 6.27, *SD* = 4.91) conditions, *t*(48) = 1.66, *p* = .10, *d* = 0.25.

### Individual Differences

The information provided by the compass may have been difficult to integrate and use as a navigational aid, except by navigators who were adept at using the information. We correlated SBSOD and SCNEQ scores with the difference in pointing error with the compass minus without the compass on each type of pointing judgment, for each session. We predicted that better navigators would have greater improvement between compass and non-compass pointing, resulting in a negative correlation. However, none of the correlations reached significance (all *r*'s < |.25|). Improvement on the external landmark pointing judgments, specifically, appeared to be common, regardless of individual differences in navigation abilities.

As an exploratory analysis, we coded participants' responses to the post-test question about whether they found the compass useful and the reasons they found it useful. (This analysis was performed by authors: SW and DB). We used a consensus process whereby each coder separately rated the response as Useful, Not Useful, or Mixed. Mixed were a heterogeneous group, consisting of participants who responded that the compass A) neither helped nor hurt them, or B) helped them in some ways but hurt them in others. Disagreements (on 7 participants' responses) were discussed and resolved. Each coder also rated what specific tasks the participant mentioned certain the compass being useful for, if any. We coded 6 participants' responses as being Not Useful, 16 as Mixed, and 26 as Useful. Of those who found the compass Mixed or Useful, 14 reported finding the compass useful specifically for Internal Landmarks. We did not observe any significant differences in performance as predicted by their self-assessment of the usefulness of the compass. Interestingly, we also did not observe any differences in self-reported sense of direction within each of these groups. Nor did we find that improvement between the non-compass and compass routes correlated with sense of direction or differed among these three groups.

## Discussion

In this study, we assessed the use of a vibrotactile compass on real-world navigation. Our aim was to determine whether providing a global directional cue improved navigational performance generally across spatial tasks, or specifically on tasks for which the global direction was directly relevant. Our data support the specific hypothesis. We found selective improvement for pointing to external landmarks, which were situated in the larger urban area with which participants were already familiar. We did not find improvement on spatial knowledge of internal landmarks, either measured through pointing or with sketch maps. We also did not find better route learning or changes in use of spatial language as a function of wearing the compass. Finally, we found that people used the compass more effectively to point to external landmarks because those landmarks were coded with respect to global north. These data suggest that vibrotactile compasses may help integrate location in a new space with a familiar, wider reference frame.

These data support the specific hypothesis – providing global directional cues resulted in selective improvement for navigational tasks that could be solved by knowing the location of true north. The best fit analysis revealed that despite being disoriented with respect to north, participants' internal configuration of external landmark directions was retained; the reference frame in which the external landmarks were encoded – north – was merely rotated. The compass anchored this reference frame, reducing pointing error because the configuration of external landmark judgments was already coded in an internally consistent reference frame. On the other hand, the best fit analysis revealed that people did not encode the internal landmarks using the same reference frame as external landmarks. Rotating the internal landmarks did not reduce pointing error; nor did it yield a best fit angle that correlated with where the participant thought north was. This observation suggests that participants in this study did not encode the internal landmark orientations or locations with respect to global north. Providing participants with a directional cue did not prompt them to use a place-based strategy for learning the locations of landmarks or the routes. It also did not promote the use of a global reference frame to encode the indoor space, but rather provided access to the global reference frame in a space that was otherwise devoid of that information.

Since the participant only needs to pay attention to the device when pointing to external landmarks, we do not know (beyond extrapolation from self-reports) whether and how many participants ignored the device while walking or when pausing at landmarks. One alternative possibility, which the present experiment cannot rule out conclusively, is that, rather than provide access to a reference direction, the vibrotactile compass allows users to update their spatial position and facing direction during its use (Loomis, Klatzky, Golledge, & Philbeck, 1999). We attempted to test this possibility in the current study by choosing campus landmarks that changed their position relative to the navigator from the beginning to the end of one route. Unfortunately, because this change applied to only one trial per subject, we were limited in our power to observe differences. Future experiments could use optimally designed environments to test this question. Navigators could learn a novel virtual environment with landmarks that are either rendered at infinity (and thus are always in the same direction, regardless of where the navigator is in the environment) or rendered at some nearby distance so that they change direction depending on which part of the environment the navigator is currently located. If access to the spatial direction helps performance, but participants did not use the compass to update their position, then infinitely rendered landmarks should improve with the compass, but nearby landmarks would not. If navigators update their position using the compass, the ability to point to all landmarks should improve.

The dissociation between reference frames and navigational strategies implies distinct involvement of different brain systems – the hippocampal-based place learning system and the caudate-based route learning system. Research in humans (Hartley, Maguire, Spiers, & Burgess, 2003; Marchette, Bakker, & Shelton, 2011) and rodents (McDonald & White, 1994; Morris, Garrud, Rawlins, & O’Keefe, 1982; Munn, 1950; Packard & McGaugh, 1996; Restle, 1957; Tolman, Ritchie, & Kalish, 1946) has shown that recalling a sequence of turns – route learning – depends on the caudate, whereas learning locations and directions in an environment – place learning – depends on the hippocampus. Our results suggest that the hippocampal place learning system is involved when navigators incorporate compass direction into an existing representation of the external environment. Because the external landmarks and the newly-learned internal landmarks had never been travelled between directly, the directions between them (i.e., novel shortcuts) had to be inferred. Such an inference could only be made with place learning. On the other hand, the compass cues did not aid the route-learning system, likely responsible for recalling the sequence of turns in the hospital, either because navigators did not need it (i.e., the route was easily recalled without it), or because it was not deemed helpful. Why learning the internal landmark locations was not easier with the compass suggests that the place learning system was either less involved in learning the layout of the hospital, or this information was not directly helpful to the route learning system. Either of these possibilities, as well as confirming the involvement of different brain regions for the different navigational tasks awaits future study.

In the present study, participants were not disoriented in the indoor environment. Overall, they could reverse the route making very few errors, with or without the compass. This ability suggests a dissociation not just between internal and external reference frames, but a dissociation between how different navigation tasks might rely upon distinct reference frames. Route reversal requires recalling a sequence of turns and intersections, but does not require mapping the configuration of the route onto a global reference frame. In other words, participants could remember the order of four turns (e.g., left, left, right, left), and thus recreate the route without knowing whether they were facing the same direction at the beginning and the end. One possibility is that the compass would have helped participants learn more complex routes than those tested here. Another possibility is that the compass might help people with memory deficits like Alzheimer’s Disease, who might not be able to retain a memory of the navigational choices made in a learning a new route. These possible uses of the compass await future research.

Unlike route-reversal, pointing judgments require a consistent reference frame on which the configuration of the route must be situated. Despite accessing a global reference direction provided by the compass, this information did not improve performance for internal landmarks. The external environments were familiar to our participants, and are frequently viewed on maps, or in ways that demand knowledge about their global positions (e.g., which direction to drive, walk, or take a subway line to). Because the external environments are already linked to this global reference frame, providing information about north allows navigators to orient to landmarks they have coded with respect to north. In the unfamiliar indoor environment, placing those landmarks in a global map relies on path integration – the ability to maintain and track direction and translation through the space. These findings relate to azimuthal reference, which is an anchoring point in the environment that provides a heading for a navigator. The global reference direction provided by the vibrotactile compass – north – allows heading to be directly sensed in the environment (Loomis, Klatzky, Golledge, & Philbeck, 1999), in a way that is not possible with cues from the internal environment alone. Importantly, an azimuthal reference can correct path integration, but successful path integration does not need an azimuthal reference. One can combine inertial cues with rotational cues from head direction cells (Ranck, 1985; Taube, Muller, & Ranck, 1990) to calculate a reference direction without reference to an azimuth. That is, a navigator may be oriented within a local environment, but not know how to link the local environment orientation to the global environment. Providing a global reference direction aids one aspect of this process – navigators know the direction they are travelling with respect to the global environment – but they have no information about distance or the azimuth. Without information about distance, recreating the route is still possible (i.e., using a response-based strategy), but attaining metric knowledge about the spatial configuration of landmarks is not.

We do not know if a different testing environment would yield different results for either internal landmark accuracy or knowledge of the route configuration. Two aspects of the hospital environment used here may have predisposed participants to rely on response-based information when building route knowledge: 1) a highly-structured environment, consisting mainly of 90° turns, and 2) a relatively large number of such turns. These properties could mean that participants easily kept track of their rotation after each individual turn, but they might not have integrated the large number of turns into one representation. This possibility leads to the prediction that the vibrotactile compass might be useful in situations where a navigators have difficulty updating the effects of rotations because of non-canonical turns, and if taking many turns. Testing for the usefulness of a compass in updating internal landmarks in controlled virtual environments, where such properties can be varied while others can be kept constant would provide a better test of these hypotheses.

In a sighted population, vibrotactile north did not promote more accurate general spatial knowledge. As previous work has shown and as we report here, participants easily report the direction being indicated by the compass, showing little difficulty incorporating an embodied and previously unfamiliar cue of an abstract property into concrete spatial information (Kärcher et al., 2012; König et al., 2016). Nevertheless, providing global north is not a panacea for improved navigation. Navigation is multifarious. Multiple strategies are used to learn spaces; different spatial tasks require different spatial knowledge. We outlined two ways the compass could be used to augment spatial performance – by shifting a response-based to place-based strategy, or by shifting the reference frame in which the environment was learned. The present data suggest that neither occurred. This observation emphasizes the difficulty of shifting a preferred navigation strategy or reference frame, and presents a challenge for introducing such devices for general application. Participants remained oriented with respect to the external environment, but did not successfully use this information to construct the internal environment. Moreover, participants did not use more absolute language compared to relative language when wearing the compass to describe the routes. Although language use is an indirect reflection of the nature of a spatial representation, our results suggest no difference in thought processes used by participants using different reference frames in describing the route directions.

Several limitations of this study warrant future research. First, although this population did not use a directional cue to enhance internal landmark knowledge, we do not know if the result would generalize to compromised populations. Blind people might be able to use directional information more effectively to learn a new space, since they can effectively integrate paths without visual information (Loomis et al., 1993). As mentioned earlier, we also do not know if the directional information would help people with memory deficits of the kind that occurs in Alzheimer’s disease.

Second, the vibrotactile compass intervention might not have overcome the variability of individual differences. It is possible that good navigators did not improve with the compass; bad navigators could not use the information provided by the compass; but the intermediate group did improve. We looked for, but did not find such effects. The strategy questionnaire did suggest individual differences in compass use – some participants liked using the compass, while others tried their best to ignore it. These differences in compass use did not relate to differences in performance, however. Relatedly, the hospital's axes were misaligned by approximately 9° with the cardinal directions. The geometry of environments is a powerful cue and can drive the choice of preferred reference frames (Kelly, McNamara, Bodenheimer, Carr, & Rieser, 2008; Shelton & McNamara, 2001). This preference may have led fewer participants to use the compass than otherwise would have if the cardinal directions and building axes were aligned.

Third, our intervention was minimal. Although we instructed participants on how the compass worked, we did not suggest spatial strategies (e.g., try mapping the new environment using the compass direction). Nevertheless, we observed large effects of the vibrotactile compass on global orientation. These results suggest that these devices could be effective with minimal training and brief exposure. Past research with vibrotactile compasses has revealed changes in spatial language and behavior over longer time spans (weeks; Kaspar, König, Schwandt, & König, 2014), but not in the small dose and short duration we used. In those studies, participants wore the device every day, and reported a more natural, integrated sensory experience with the vibrotactile compass. More direct instruction or more experience with the vibrotactile compass might improve additional aspects of navigation behavior. Some participants reported that they ignored the compass, or were confused by it. They may have gotten used to the device if used longer than an hour. In a more controlled setting, participants could be coached to incorporate directional information through increasingly complex routes, being required to point back to start after certain intervals. In this way, a reference direction might augment path integration. More extensive training might also ameliorate individual differences. For example, the extent to which individual participants use visual compasses in their everyday navigation experience could affect the degree to which they adopt an alternative means of accessing that information. Further training for individuals who do not readily use global directions to navigate might benefit them.

Finally, the type of selective improvement provided by knowledge of global information may be affected by the modality in which the information is provided. Since we were interested in the extent to which providing a global reference direction influenced aspects of navigation behavior, we did not vary the kind of sensory information that provided the reference direction. For example, alternative conditions could have used standard visual compasses, or provided continuous verbal feedback as the participant navigated. Ultimately, we predict our findings would generalize to alternative input modalities, as long as participants are not distracted by looking down at a compass or listening to information while walking. This prediction is because we found that participants did access the global direction with the vibrotactile compass, but did not necessarily use it to update their representation of internal landmarks, and so should be accessible similarly through different sensory modalities. Still, offering auditory or visual cues of global direction either continually or at each landmark might give a form of information that is more easily integrated with internal reference frames.

In sum, we found limited utility of a vibrotactile compass for learning a new indoor environment. These findings further support cognitive research on the use preferred reference frames for encoding spaces. The present results advance this work to show that new environments are not mapped onto familiar reference frames, like those anchored to north. The results also emphasize the importance of decomposing aspects of navigation behavior to determine not just whether navigation improves, but how. Finally, future work seeking to improve navigation behavior with compasses should consider that the benefits of vibrotactile compass use for indoor navigation are likely to stem from mapping a new space onto a familiar one, not from learning a new space in a different way.

## Acknowledgments

We would like to thank Ryan Boswell for assistance in developing the custom-built pointing device, and mending the vibrotactile compass.

## Footnote

1 For two subjects, we used a custom built Arduino device which participants could hold in their hand, point in the direction, and press a button to indicate the direction. While this method had ergonomic benefits, and ensured that participants could not accidentally view the digital compass, calibration with the digital compass revealed that this method yielded less reliable measurements.

2 Internal landmark judgments were combined so that the total number of pointing judgments used in the best fit analysis was more closely matched to the number of external landmarks (eight for external landmarks, six for internal landmarks). The start/end pointing judgments were excluded because these were not equated across time points. Simulations with random data reveal that fewer landmarks are easiest to fit with a best-fit analysis. With random data, the average error of three pointing judgments, after best-fit rotation, is approximately 45°; 6 pointing judgments have approximately 62°; 8 pointing judgments have approximately 66°. We determined that because error could be reduced more effectively with fewer judgments, that the number of judgments between internal and external landmarks should be as close as possible. Nevertheless we find the same results when location and orientation judgments are analyzed separately. This decision introduces the possibility that different best-fit solutions might exist for the two sets internal landmark judgments (orientations versus locations). However, in the current data, the best-fit rotations obtained for the locations and orientations separately were correlated highly (circular correlation r = .45, p = .006).

